# The economy and environmental attitudes: Does a good economy make citizens care more about the environment?

**DOI:** 10.1101/502682

**Authors:** Stephanie M. Rizio, Yoshihisa Kashima

**Affiliations:** Melbourne School of Psychological Sciences, The University of Melbourne, Parkville, Victoria, Australia.

**Keywords:** environmental attitudes, GDP, economy, perception of economy

## Abstract

The relationship between economic conditions and environmental attitudes has been hotly debated in academic and public discourse. Some contend that economic prosperity strengthens environmental attitudes, while others argue that no such relationship exists. We shed light on the economy-environmental importance relationship by incorporating both nonlinear effects and public perceptions of the economy. Two countries with comparable economic performance and cultural heritage, Australia and Canada, are investigated. Using opinion survey data in a time-series analysis over a 20-year period, we find that GDP per capita has a nonlinear relationship with environmental attitudes in both countries, and the inclusion of perceptions of the economy significantly improves the prediction of environmental attitudes. In both Australia and Canada, a U-shaped relationship is observed; initial improvements in economic conditions tend to worsen environmental attitudes, but past a certain threshold, further economic improvements result in increased environmental concern. Perceptions of the economy too showed analogous trends. In both countries, positive perceptions of the economy were associated with strong environmental attitudes; at least in Canada, negative perceptions of the economy were also positively associated with environmental attitudes. The nonlinear effects of the economy as well as perceptions of the economy helps shed light on the current inconclusive literature on the relationship between the economy and citizens’ pro-environmental support. This is an important consideration given the critical importance to the future sustainability of the natural environment.

---

“It’s the economy, stupid!”

*James Carville, former campaign strategist to US President Bill Clinton*

## Introduction

Environmental sustainability and the state of the natural environment is one of the most significant issues in the world today (Intergovernmental Panel on Climate Change, 2014; McMichael et al., 2004). Humankind’s attitudes and actions are of critical importance to the future sustainability of the natural environment. One of the most touted, but also contentious, determinants of the public attitudes towards the environment is the economy, namely, a human-made system consisting of the production, distribution, trade, and consumption of limited goods and services by different agents in a given geographical location (Bernanke, 2004). Generally, the state of the national economy is seen to be a robust predictor of a nation’s environmental importance (e.g. Kahn & Kotchen, 2010; Scruggs & Benegal, 2012; Shum, 2012), as broadly defined as environmental attitudes which include a genuine concern for the natural environment, as well as attitudes towards engaging in environmental degradation-avoidance practices, such as recycling or curbing electricity use (e.g., Steg & de Groot, 2012; Mayer & Frantz, 2004; Blake, 2001; Stern, 2000; Stern, Dietz & Guagnano, 1995).

Indeed, there is substantial evidence that, over time and across countries, people’s attitudes towards the environment vary as a function of the state of the economy (e.g., Bolsen & Cook, 2008; Brulle, Carmichael, & Jenkins, 2012; Harring, Jagers, & Martinsson, 2011; Kachi, Bernauer & Gampfer 2015; Kahn & Kotchen, 2010, Pew Research, 2009; Scruggs & Benegal, 2012). However, there is much debate about the direction of the relationship. Some have theoretically argued for or empirically shown a *positive* relationship between the economy and environmental attitudes. That is, a wealthy national economy is associated with increased environmental importance (i.e., environmental attitudes and sustainability, as well as willingness to protect the natural environment; Inglehart 1990, 1995, 1997; Diekmann & Franzen, 1999; Franzen, 2003; Franzen & Meyer, 2010; Kahn & Kotchen, 2010; Scruggs & Benegal, 2012; Conroy & Emerson, 2014), whereas others have contended the reverse (i.e., the stronger the economy, the lower the environmental attitudes; Dunlap & Mertig, 1995, 1997; Gelissen, 2007). Some other studies find no relationship between the economy and environmental attitudes (Baek, 2015). Thus, the question remains: how is the state of the economy related to perceived environmental importance? The present article examines this question in Australia and Canada over 20 years.

### How is the Economy Related to Environmental Attitudes?

A variety of approaches have been used to investigate the economy-environmental importance relationship in the past. First, investigating the relationship between objective indicators of the economy (e.g., GDP per capita) and environmental attitudes across countries, a number of studies found that richer countries are more concerned about and make greater efforts to protect the natural environment (Conroy & Emerson, 2014; Franzen & Meyer, 2010; Grossman and Krueger, 1991; Kahn & Kotchen, 2010; Alibeli & Johnson, 2009; Owens & Videras, 2006; Scruggs & Benegal, 2012; Turaga, 2015). However, there are some indications that poorer countries also are equally or even more concerned about their environment (Dunlap & Mertig, 1995; Gelissen, 2007). Nonetheless, various country-level confounds may hamper a clear determination of the direction of the relationship between the economy and environmental attitudes in these studies. For instance, the actual state of the natural environment, including the levels of air, water, and land pollutions, may be a more important determinant in this case (e.g., Dunlap & Mertig, 1995).

Second, some studies investigated the relationship within a country over time (Conroy & Emerson, 2014; Kahn & Kotchen, 2010; Scruggs & Benegal, 2012; Turaga, 2015). With the exception of Conroy and Emerson’s 40-year analysis, these studies typically used the nation’s objective economic indicators for a short period of time, generally showing that as the country’s economic indicators improve, environmental attitudes grow, i.e., a positive relationship. However, there are some indications that the relationship is nonlinear. The notion of the Environmental Kuznets Curve (EKC) (Grossman & Krueger, 1991) suggests that when the economy is poor, objective economic conditions may negatively relate to environmental attitudes, but as the economy reaches a certain level of prosperity, objective economic conditions may positively relate to environmental attitudes. This is because a nation can afford to dedicate economic resources to environmental sustainability only when a certain level of wealth is attained (cf. Inglehart, 1974). Furthermore, at even higher levels of economic wealth, the importance placed on environmental issues may start to wane because people believe that they have fulfilled their obligations for environmental protection, for instance, by paying environmental taxes (Meijers, Verlegh, Noordeweir, & Smit, 2015; Tiefenbeck, Staake, Roth, & Sachs, 2013).

In accordance with this line of thinking, Baek (2015) analysed nonlinear trends across Arctic circle nations and found that GDP per capita had a negative impact on CO_2_ emissions in Canada, Denmark and the U.S, but had no significant effect in Finland, Iceland, Norway and Sweden. In addition, Baek (2015) documented that the relationship between the economy and environmental attitudes was nonlinear in Sweden. Thus, the theoretical reasoning and empirical findings for nonlinearity highlights the complexity of the relationship between the economy and environmental attitudes. It suggests that the relationship may be positive or negative depending on the range of economic wealth that a study examines.

Finally, the studies cited above did not investigate people’s *perceptions* of the economy and their relationship with environmental attitudes. Those that did (Durr 1993; Elliott, Seldon, & Regens, 1997; Harring et al., 2011, Kachi et al., 2015), showed that their relationship to environmental attitudes is not straightforward. For example, in Sweden, the relationship depended on the observation period (Harring et al., 2011): prior to the change in government and economic policy of 1997, there was a positive relationship, but after then, the correlation between the perceived economy and environmental attitudes was no longer seen. Such complexities may be due to the discrepancy between the objective and subjective evaluations of the economy. Indeed, a number of studies have shown that factors such as political affiliation, global “mood” or sociotropic attitudes, media, and peer effects may bias people’s perceptions of the economy (McCright & Dunlap, 2011; Brulle et al., 2012; Cragg, Zhou, Gurney, & Kahn, 2013; Atcherberg, 2006; van de Bles, Postmes & Meijer, 2015; Ansolabehere, Meredith, & Snowberg, 2014; Millner & Ollivier, 2016; Goeschl, Kettner, Lohse, & Schwieren, 2013; Heath & Gifford, 2006).

### Current Study

In the current study, we investigate the relationship between (i) objective economic conditions and perceptions of the economic conditions with (ii) environmental attitudes over time and within two countries: Australia and Canada. They were selected because of their similarity in cultural heritage, economic composition and size. We will examine (1) whether there is a nonlinear relationship between objective economic conditions and environmental attitudes, and (2) whether popular perceptions of the economy contribute to the prediction of the importance of environmental issues. Objective economy is measured by GDP per capita, whereas perceptions of the economy is measured by (a) the proportion of survey respondents who believe the economy has improved (% positive), and (b) the proportion of those who believe the economy has deteriorated (% negative) because different segments of the population may form different perceptions of the economy (Santos, 1970, p. 234). We do not include % neutral in our regressions, since % neutral is a linear combination of % positive and % negative, which would result in multicollinearity (% neutral = 100 - % positive - % negative). We perform a time-series analysis covering more than twenty years. This is a significant length of time, making it possible to investigate how long-run economic developments and business cycles affect the relationship between the economy and environmental attitudes. In macroeconomics, the long run is a period where the general price level, wage rates, and economic expectations adjust fully to the state of the economy, whereas the short run is when these variables may not fully adjust (Samuelson & Nordhaus, 2004). Business cycles, also a macroeconomic phenomenon, occur due to disturbances to the economy that push economic activity into phases that are above or below its full level employment (Romer, 2008).

## Method

Australia and Canada are selected based on their relatively similar economic profile: the two countries are similar in economic development and labor force composition. In 2014, Australia and Canada had scores of 0.935 and 0.913 out of 1 on the United Nations Human Development Index (HDI; whose component variables relate to health, education and standard of living measured as a geometric mean of normalized indices for each of the three dimensions), a summary measure of average achievement in key dimensions of human development, ranking 2^nd^ and 9^th^ respectively. In 2014, the two countries also experienced similar rates of economic growth (2.7% for Australia and 2.4% for Canada). The economic composition in both countries were also similar in 2014: agriculture ranged from 1-3% of Gross Value Added (GVA; a measure of the contribution to GDP made by an individual producer, industry or sector calculated by the value of output less the value of intermediate consumption, OECD, 2011), industry from 27 per cent to 31 per cent of GVA, and services and other activities ranged from 69 to 70 per cent of GVA (United Nations, 2016).

### Data Sources

Over 20 years of data are captured for Australia and Canada, covering years 1990 to 2013, for a time-series analysis. Following the convention in the social sciences literature, we employ GDP per capita at current international prices, adjusted for Purchasing Power Parity (PPP) (World Bank, 2011), as our objective indicator of economic performance.

The survey data used for each country are as follows (see also Table 1):

**Table 1.** **Survey items used to measure variables – Australia and Canada.**

| Variable | Source | Years surveyed | No. observations | Survey item | Survey item wording | Response scale |
| --- | --- | --- | --- | --- | --- | --- |
| *Australia* |  |  |  |  |  |  |
| Environmental attitudes | ANU<br>http://www.australianelectionstudy.org/ | 1990 1993, 1996, 1998, 2001, 2004, 2007, 2010, 2013 | 1,769 to 3,955 (averaging 2,411) | Most important election issues | 1990: ‘Which of these issues has worried you and your family most in the last 12 months?’<br>1993: ‘Still thinking about the same 14 issues, which of these has worried you and your family most in the last 12 months?’<br>1996: ‘Still thinking about these 13 issues, which of these issues has been most important to you and your family?’<br>1998-2013: ‘Still thinking about these 13 (2007: 14, 2010: 12, 2013: 10) issues, which of these issues has been most important to you and your family during the election campaign?’ | Selected from a provided list of (10 to 14) items |
| Subjective economy | ANU<br>http://www.australianelectionstudy.org/ | 1990, 1993, 1996, 1998, 2001, 2004, 2007, 2010, 2013 | 1,769 to 3,955 (averaging 2,411) | Financial situation of country over past year | “What do you think has been the financial situation of Australia over past year?” | 1 Become better<br>2 Become worse<br>3 About the same |
| *Canada* |  |  |  |  |  |  |
| Environmental attitudes | CORA<br>http://www.queensu.ca/cora/trends/ | 1990-2011 (except 2003) | 1,500 and 2,084 (averaging 1,986) | Most important problem in Canada | In your opinion, what is the most important problem facing Canadians today?” | Selected from a provided list of 30 items |
| Subjective economy | CORA<br>http://www.queensu.ca/cora/trends/EC_General.htm | 1988, 1993, 1997, 2000, 2004, 2006, 2008 and 2011 | 869 and 1,576 (averaging 1,375) | Economy over past 12 months | In your opinion, is the Canadian economy getting stronger, weaker or is it staying about the same? | 1 Get better<br>2 Stay same<br>3 Get worse |

#### Australia

The Australian National University (ANU) national post-election survey covering years 1990, 1993, 1996, 1998, 2001, 2004, 2007, 2010 and 2013 was used to measure environmental attitudes and the perceptions economy. Respondents to the survey were drawn randomly from electoral registers. The number of respondents to the survey question ranged from 1,769 to 3,955 (averaging 2,411).

#### Canada

Data on Canadian environmental attitudes were obtained from The Environics Institute for Survey Research’s Focus Canada survey, which was conducted every year from 1990 to 2011 (except 2003). The number of respondents ranged from 1,500 to 2,084 (averaging 1,986). Data on the subjective economy were obtained from the Canadian National Election Survey (CNES), conducted in years 1988, 1993, 1997, 2000, 2004, 2006, 2008 and 2011. The number of respondents to the survey question ranged from 869 and 1,576 (averaging 1,375). Both data sources were accessed from the Canadian Opinion Research Archive (CORA) at Queen’s University.

### Measurement Construction

In measuring the objective economy, we used a natural logarithmic scale to measure the effect of GDP per capita more clearly. To contextualize our findings, however, the scaled GDP per capita values were converted back into US dollars (2011 price level) when reporting on results. Perceptions of the economy (subjective economy) were measured as the proportion of the respondents who believed that the economy had gotten better (% positive) over the past 12 months and the proportion of the respondents who believed that the economy had gotten worse (% negative).

Environmental attitudes were measured by the proportion of the respondents who selected the environment in response to the question, “What is the most important problem [in your country]”. The number of options given to this question varied across countries and years. Therefore, an adjustment for the variation in the number of options given to a respondent was made, where the unadjusted value in a given year is multiplied by an adjustment factor. The adjustment factor was calculated by dividing the number of options in that year by the average number of options throughout the entire period in which survey data were available.

Where data could not be obtained, the values for each variable were interpolated for years in the period investigated. The interpolation of missing data was done by averaging the values before and after the missing value. For example, to interpolate for a year, t, where data for the previous year (t-1) and succeeding year (t+1) exist, the year t value will equal the average of values for years t-1 and t+1. If there was more than one year between observed values, the same interpolation is made where a constant growth rate is assumed from one year to next. Interpolated data are often used in time series analysis because observed and interpolated values are typically well correlated (Friedman, 1962, p. 731).

## Models

The time series models are specified as follows:

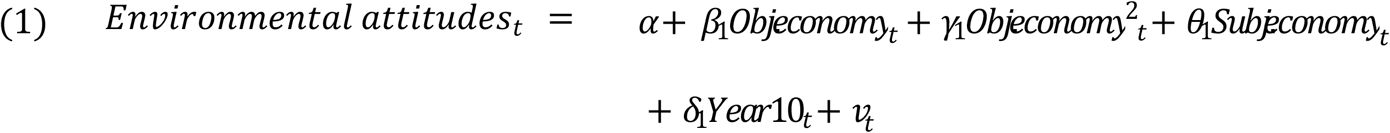

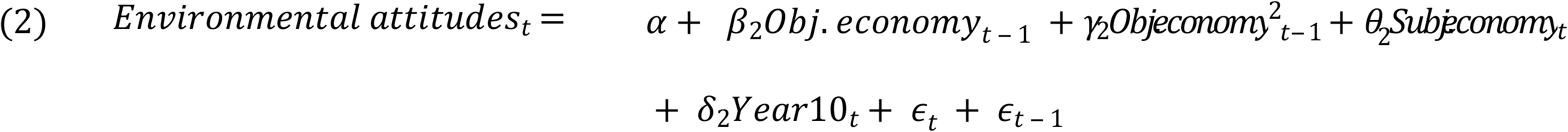

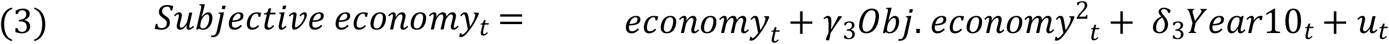

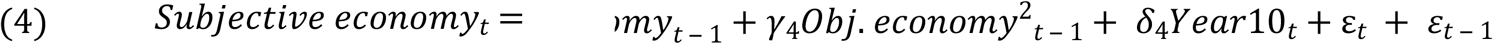

Equations (1) and (2) test the effect of objective economic conditions and perceived economic conditions on environmental attitudes, and equations (3) and (4) test whether the subjective economy can be predicted by the objective economy. A lag of one year is included for the objective economy (including the error term) in equations (2) and (4) to account for the time it may take for the objective state of the economy to be recognized by respondents. The subjective economy variable is not lagged as perceptions of the economy are likely to affect environmental attitudes in real-time, rather than with a delayed adjustment. All equations include a constant term (α), and an error term (ε), which measures the variation not explained by the included independent variables.

We include (i) quadratic and linear terms for the objective economy to test for a nonlinear relationship; and (ii) a dummy variable to see if there is a change in environmental attitudes from the first ten years, compared to the period from the 11^th^ year onwards (set equal to 1 from 2000 to 2013 and 0 prior to 2000). This is because policy objectives aimed at pro-environmental support—particularly towards climate change—were less common in Australia and Canada in the first ten-year period relative to latter years, when several pro-environmental policy initiatives were introduced. Namely, the introduction of the Clean Energy Act in Australia under the Rudd-Gillard government from 2007, and the Clean Air Regulatory Agenda (CARA) in Canada under the Harper government from 2006, to decrease the level of emissions.

The Newey-West estimator was used in this analysis to correct for autocorrelation and heteroskedasticity in the error terms that may arise in a time-series. The problem with autocorrelation is often found in time-series data, where the error terms are correlated over time. Thus, the Newey-West estimator assumes that as the time between error terms increases, the correlation between them decreases (Newey & West, 1987). Finally, for equations (1) and (2), we test whether a model that includes the subjective economy provides a significantly better fit for predicting environmental attitudes than a model that excludes it.

## Results

### Australia

#### Predicting environmental attitudes

To clearly see improvements in model fit, columns 1 and 2 in Table 2 show estimates environmental attitudes (equations 1 and 2) which exclude the subjective economy measure, and column 3 in Table 2 estimates environmental attitudes by excluding the objective economy measure. Columns 4 and 5 in Table 2 show estimates of the full model (equations 1 and 2).

Columns 1 and 2 in Table 2 shows that the equation without a time lag (Equation 1) performs somewhat better, in terms of explanatory power as indicated by the adjusted R^2^, compared to the equation with time lag (Equation 2). However, when perceptions of the economy are included (columns 4 and 5 of Table 2), both equations had similar levels of fit. Therefore, for simplicity, we focus on the equation without a time lag (Equation 1). We also conducted an F-test to see whether a model that includes a measure of the perceptions of the economy provides a better fit, than a model that does not. Column 4 in Table 2 shows that the inclusion of subjective measures of the economy clearly improves on the estimation with only the indicators of objective economy (column 1 in Table 2, F(2, 18) = 10.90, p. < .01.

Looking at the parameter estimates of Equation 1 (column 1 in Table 2), the linear term is negative and the nonlinear term is positive, indicating a U-shaped relationship between the objective economy and environmental attitudes (see Figure 1). At lower levels of GDP per capita, the relationship is negative, but at higher levels, it is positive, with the inflection point at approximately $US30,000 GDP per capita (2011 price level). The Year10 dummy, is not statistically significant, indicating that there are no unobserved factors which are starkly different across the two decades sampled. The parameter estimates of Equation 1 (column 4 in Table 2) shows that positive perceptions of the economy are positively associated with environmental attitudes (see Figure 2). The greater is the proportion of the respondents who thought the economy was better, the more importance environmental issues were given.

**Figure 1.**
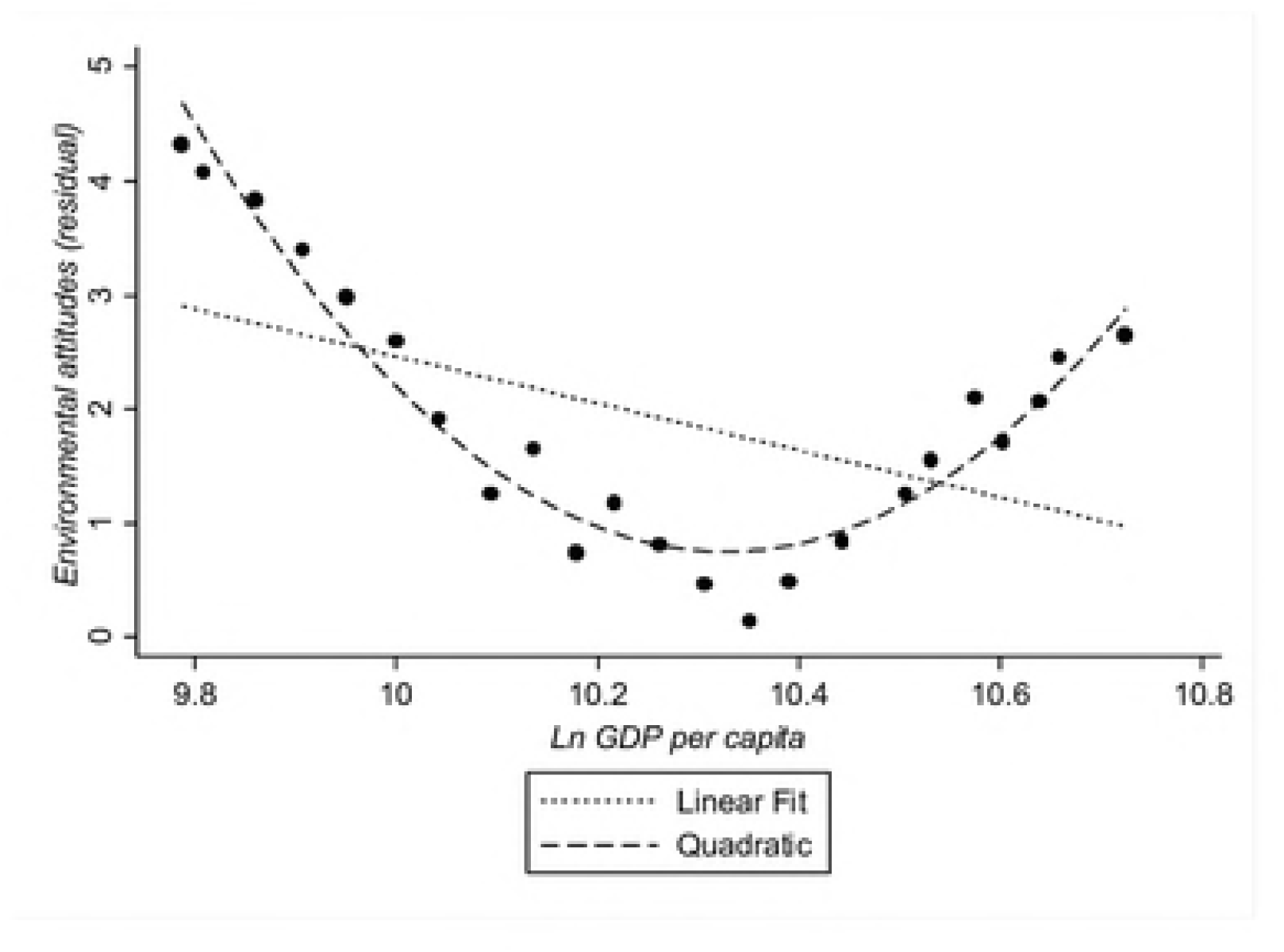
**Ln GDP per capita and environmental attitudes in Australia** *Note.* The y-axis represents the residualized value of environmental attitudes, which was computed by subtracting from the observed value, the value predicted from the estimated regression equation while holding GDP per capita at its mean, i.e., predicted value = Observed Environmental Attitudes – [1805 -.72 Year10 - .04 %negative + .13 %positive -349.2 (mean Ln GDP) + 16.91 (mean Ln GDP)^2^] (see Table 2).

**Table 2.**
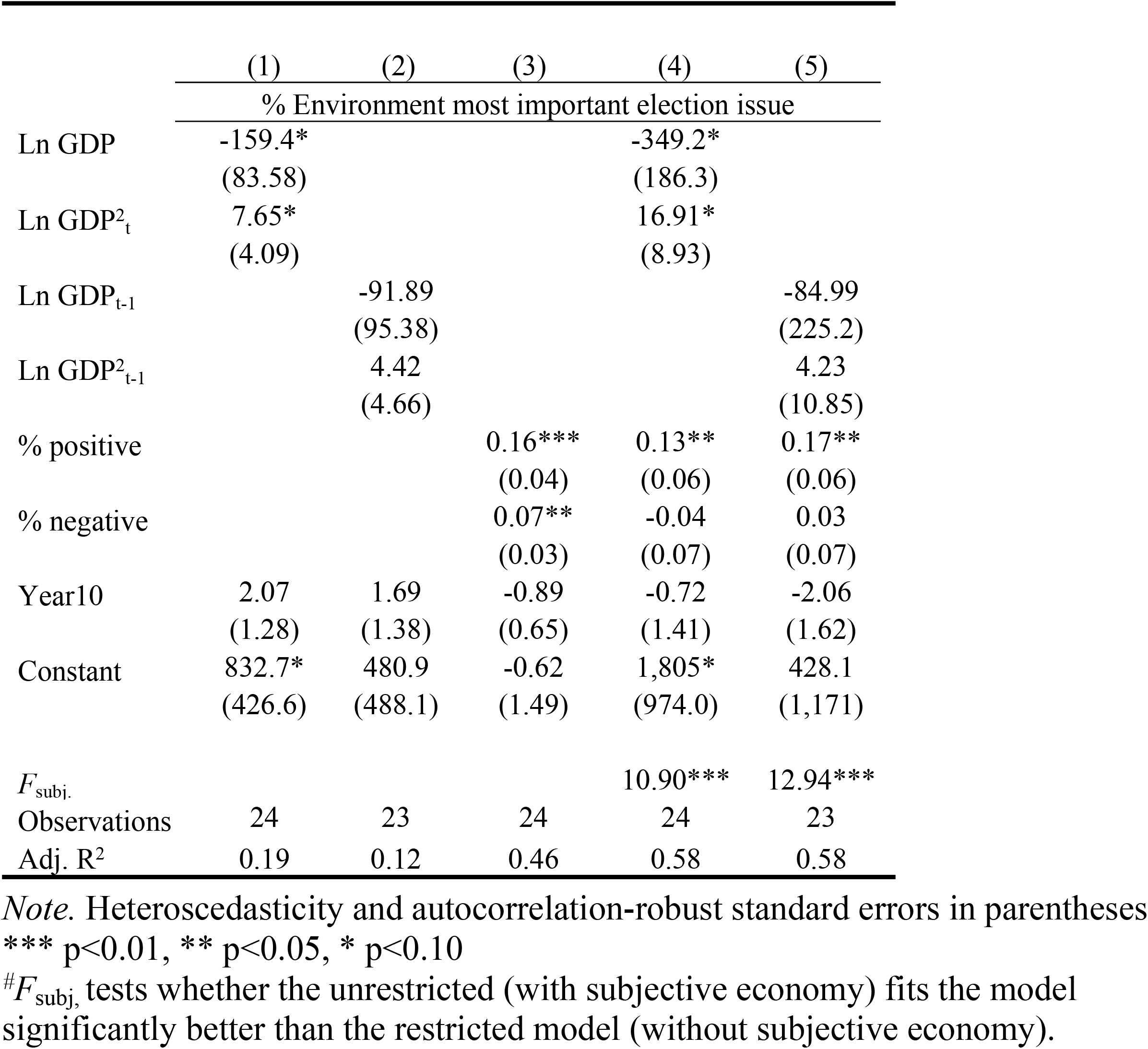
**Predicting environmental attitudes in Australia by the objective and subjective economy.**

**Figure 2.**
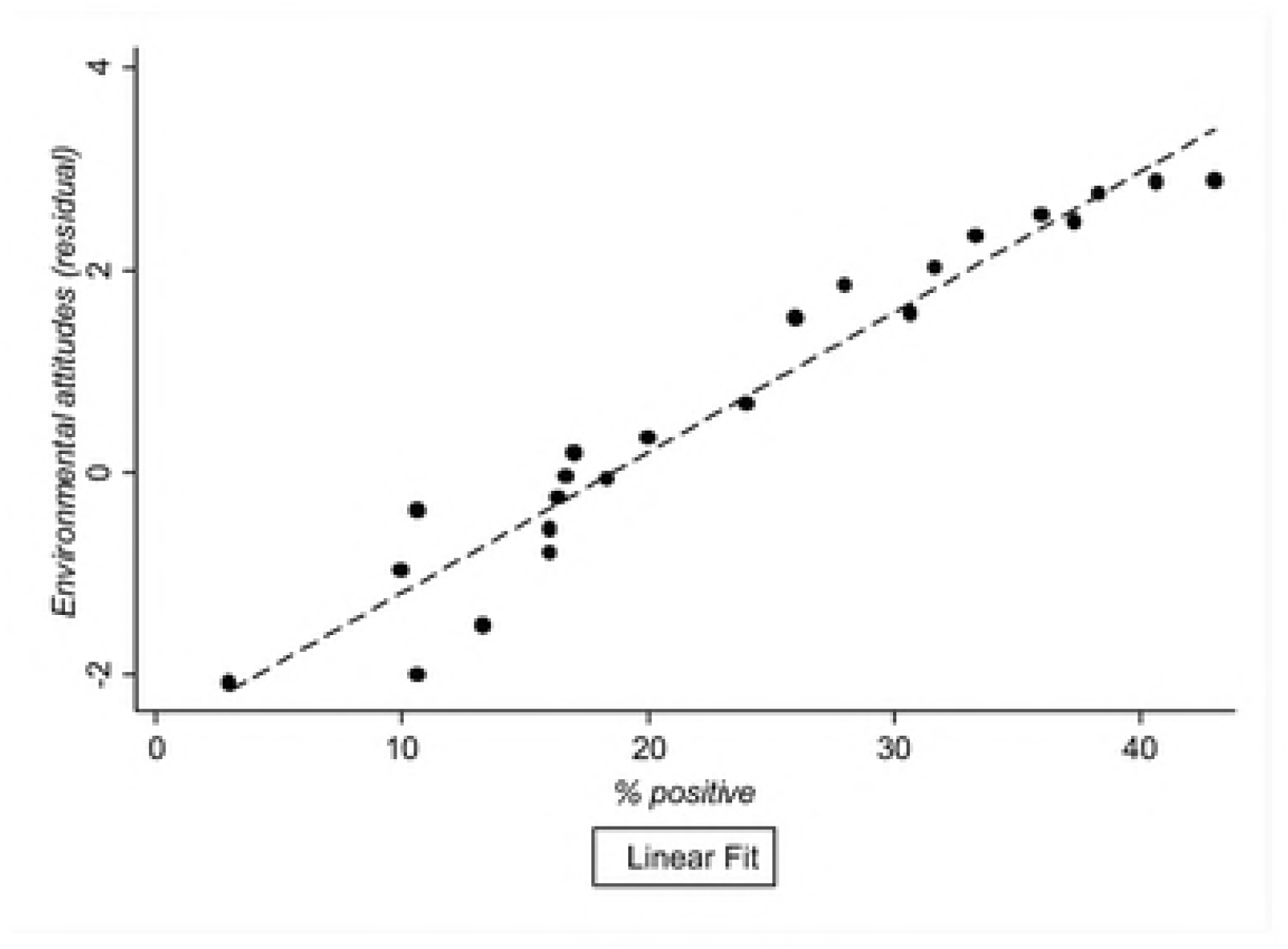
**Relationship between % positive and environmental attitudes in Australia.** *Note.* The Y-axis represents the residualized value of environmental attitudes, which was computed by subtracting from the observed value, the value predicted with the estimated regression equation while holding % positive at its mean, i.e., predicted value = Observed Environmental Attitudes – [1805 -.72 Year10 - .04 %negative + .13 (mean %positive) -349.2Ln GDP + 16.91Ln GDP]^2^ (see Table 2).

#### Predicting subjective economy

Table 3 summarizes the results for the models predicting subjective economy in Australia (Equations 3 and 4). Columns 1 and 2 in Table 3 show that positive perceptions of the economy are not significantly predicted by the objective economy. However, the only significant predictor is the Year10 dummy, suggesting that positive perceptions of the economy increased after the year 2000. In contrast, columns 3 and 4 in Table 3 show that negative perceptions of the economy are significantly related to the objective economy, explaining 91% and 90% of the variance in Equations 3 and 4 respectively. Overall, models without a time lag appear to perform better.

The nonlinear trend shows a U-shaped relationship between the objective economy and negative perceptions of the economy (Figure 3). The inflection point was at approximately $US30,000 GDP per capita (2011 price level). At lower levels in GDP per capita, increases in GDP per capita decreased negative perceptions—as one would expect. However, beyond the inflection point, an increase in GDP per capita ceased to have a strong dampening effect on negative perceptions of the economy. This implies that improvement in the objective economy does not necessarily improve economic sentiment beyond the inflection point of approximately $US30,000 in Australia. This point may be regarded as a kind of national economic reference point (e.g., Kahneman & Tversky, 1979), which its citizens expect as a satisfactory level of economic performance.

**Table 3.** **Predicting subjective economy by objective economy in Australia.**

|  | (1) | (2) | (3) | (4) |
| --- | --- | --- | --- | --- |
|  | % positive |  | % negative |  |
| Ln GDP<sub>t</sub> | 604.5<br>(424.4) |  | -2,732***<br>(259.5) |  |
| Ln GDP<sub>t</sub><sup>2</sup> | -29.95<br>(20.86) |  | 131.8***<br>(12.92) |  |
| Ln GDP<sub>t-1</sub> |  | 519.8<br>(475.6) |  | -2,959***<br>(288.8) |
| Ln GDP<sub>t-1</sub><sup>2</sup> |  | -25.98<br>(23.33) |  | 143.3***<br>(14.32) |
| 10Year | 21.70**<br>(8.81) | 22.26**<br>(9.18) | -0.57<br>(5.64) | 1.46<br>(5.55) |
| Constant | -3,037<br>(2,157) | -2,586<br>(2,423) | 14,187***<br>(1,303) | 15,307***<br>(1,456) |
| Observations | 24 | 23 | 24 | 23 |
| Adj. R<sup>2</sup> | 0.69 | 0.67 | 0.91 | 0.90 |

*Note.* Heteroscedasticity and autocorrelation-robust standard errors.
\*\*\* p<0.01, \*\* p<0.05, \* p<0.10

**Figure 3.**
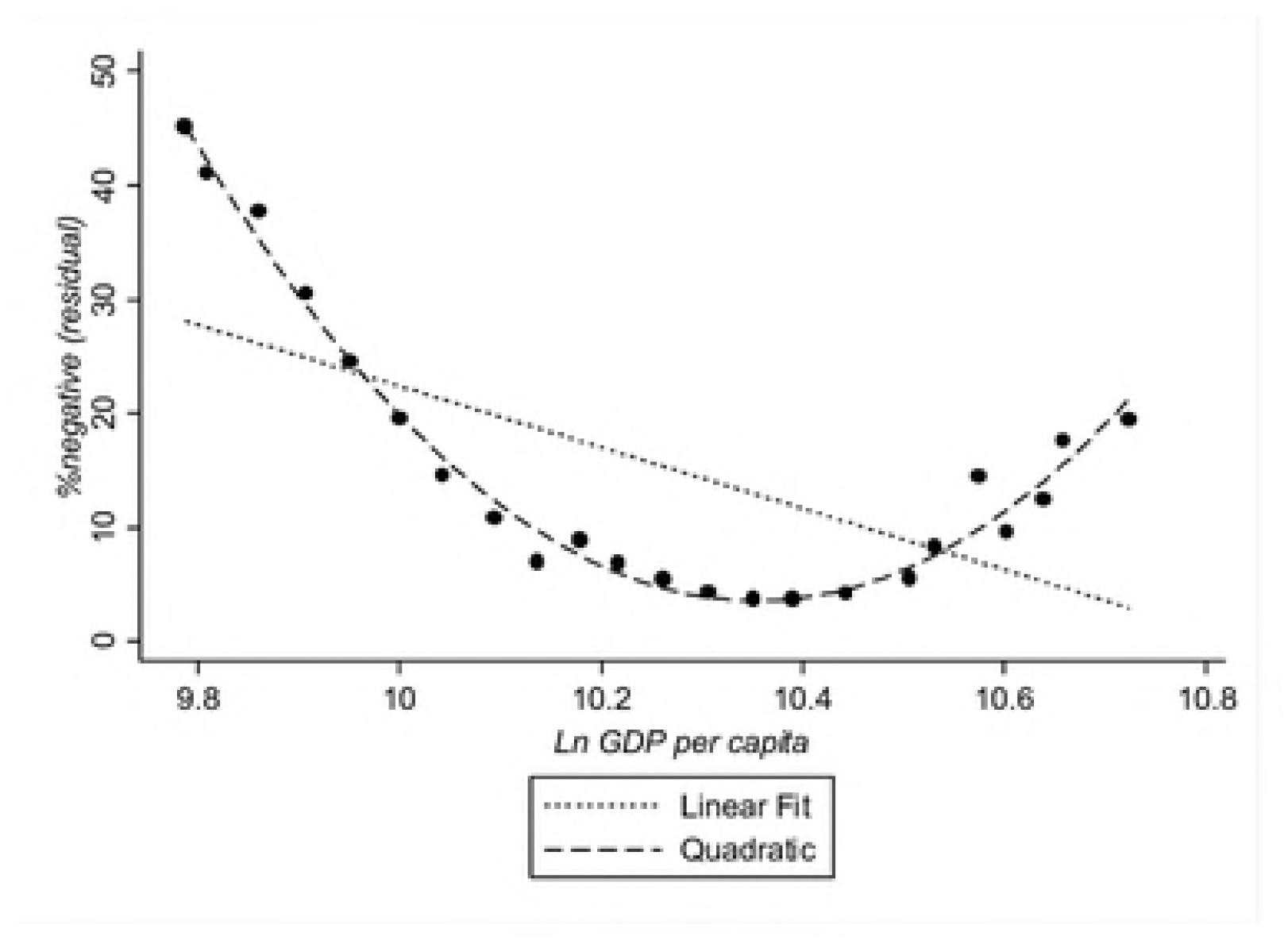
**Ln GDP per capita and % negative in Australia.** *Note*. The y-axis represents the residualized value of the proportion of the respondents who replied that the economy has gotten worse (% negatives), which was computed by subtracting from the observed value, the value predicted with the estimated regression equation while holding % negative at its mean, i.e., predicted value = Observed Environmental Attitudes – [14187 -0.57*year10 - 2732*(mean Ln GDP) + 131.8*(mean Ln GDP)2] (see Table 3).

### Canada

#### Predicting environmental attitudes

Table 4 summarizes the results for predicting environmental attitudes in Canada. Akin to the results presented for Australia (above), to clearly see improvements in model fit, columns 1 and 2 in Table 4 show estimates environmental attitudes (equations 1 and 2) which exclude the subjective economy measure, and column 3 in Table 4 estimates environmental attitudes by excluding the objective economy measure. Columns 4 and 5 in Table 4 show estimates of the full model (equations 1 and 2).

The models without and with a time lag (Equations 1 and 2) perform similarly; however, comparing predictions which include both objective and subjective economic indicators (columns 4 and 5 in Table 4), the model with a time lag (Equation 2; column 5 of Table 4) explaining 80% of the variance, does somewhat better than the model without it (71%; column 4 of Table 4). We therefore focus on Equation 2. In testing for model fit, column 5 in Table 4 shows that including the subjective economy significantly increased the variation explained (80%) compared to not including it (41%; column 2 in Table 4), F(2, 16) = 6.74, p. < .01.

Looking at the parameter estimates of Equation 4 (the best fitting model) in column 5 of Table 4, nonlinear trends for the objective economic indicators are significant, and % positive and % negative are also significant positive predictors. The negative linear and positive nonlinear terms suggest that objective economy has a significant U-shaped relationship with environmental attitudes (Figure 4). Similar to Australia, at lower levels of GDP per capita, the relationship is negative, but at higher levels, it is positive, with the inflection point at around $US27,000 GDP per capita (2011 price level). This inflection point is remarkably close to Australia’s, at $US30,000 GDP per capita (2011 price level). The Year10 dummy, is not statistically significant, indicating that there are no unobserved factors which are starkly different across the two decades sampled.

**Figure 4.**
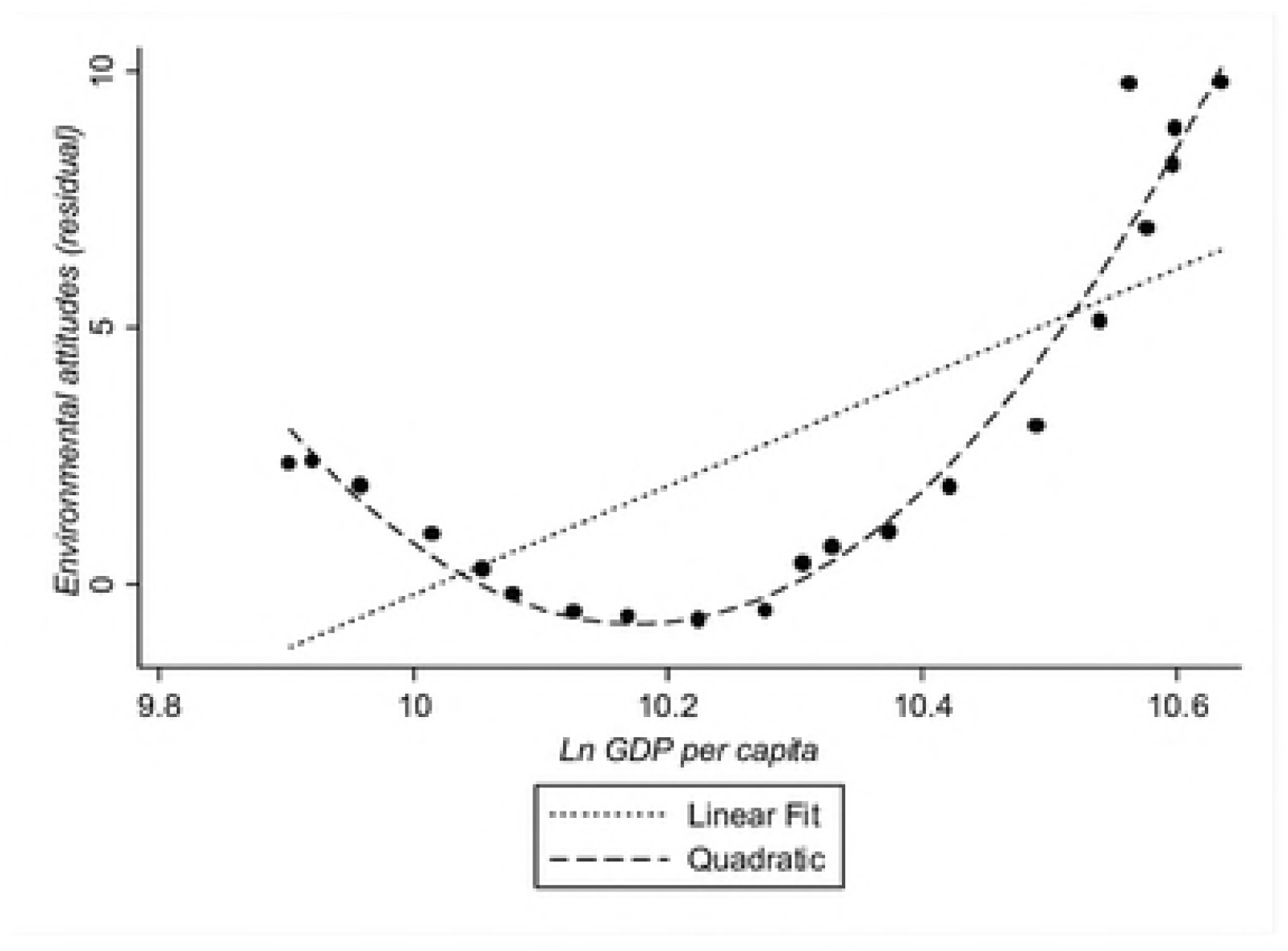
**Ln GDP per capita and environmental attitudes in Canada.** *Note.* The y-axis represents the residualized value of environmental attitudes, which was computed by subtracting from the observed value, the value predicted with the estimated regression equation while holding GDP per capita at its mean, i.e., predicted value = Observed Environmental Attitudes – [5041 + 0.37*year10 + .36*%negative + 0.61*%positive - 999.2*(mean Ln GDP) + 49.27*(mean Ln GDP)]^2^ (see Table 4).

Turning to the perceptions of the economy, both % positive and % negative were positive predictors of environmental attitudes. In particular, the greater is the proportion of the respondents who thought the economy was better, the stronger were environmental attitudes (Figure 5). In addition, the greater the proportion of those perceiving economic performance as worse, attitudes for the environment still increased (Figure 6).

**Table 4.** **Predicting environmental attitudes in Canada by the objective and subjective economy.**

|  | (1) | (2) | (3) | (4) | (5) |
| --- | --- | --- | --- | --- | --- |
|  | % Environment most important election issue |  |  |  |  |
| Ln GDP | -1,202***<br>(412.3) |  |  | -1,576***<br>(470.3) |  |
| Ln GDP<sup>2</sup><sub>t</sub> | 58.56***<br>(20.09) |  |  | 76.82***<br>(22.58) |  |
| Ln GDP<sub>t-1</sub> |  | -743.5**<br>(296.1) |  |  | -999.2***<br>(265.4) |
| Ln GDP<sup>2</sup><sub>t-1</sub> |  | 36.58**<br>(14.56) |  |  | 49.27***<br>(12.94) |
| % positive |  |  | 0.40<br>(0.31) | 0.53***<br>(0.16) | 0.61***<br>(0.16) |
| % negative |  |  | 0.33<br>(0.21) | 0.27*<br>(0.14) | 0.36***<br>(0.11) |
| Year10 | 2.11<br>(2.13) | 1.04<br>(1.50) | 4.55*<br>(2.40) | 1.68<br>(2.45) | 0.37<br>(1.62) |
| Constant | 6,165***<br>(2,114) | 3,780**<br>(1,506) | -17.55<br>(13.90) | 8,061***<br>(2,450) | 5,041***<br>(1,358) |
| $F_{\text{subj.}}$ | | | | 5.84*** | 6.74*** |
| Observations | 22 | 21 | 22 | 22 | 21 |
| Adj. R<sup>2</sup> | 0.42 | 0.41 | 0.21 | 0.71 | 0.80 |

*Note.* Heteroscedasticity and autocorrelation-robust standard errors.
\*\*\* p<0.01, \*\* p<0.05, \* p<0.1.
$\#F_{\text{subj.}}$ tests whether the unrestricted model (with subjective economy) fits the data significantly better than the restricted model (without subjective economy).

**Figure 5.**
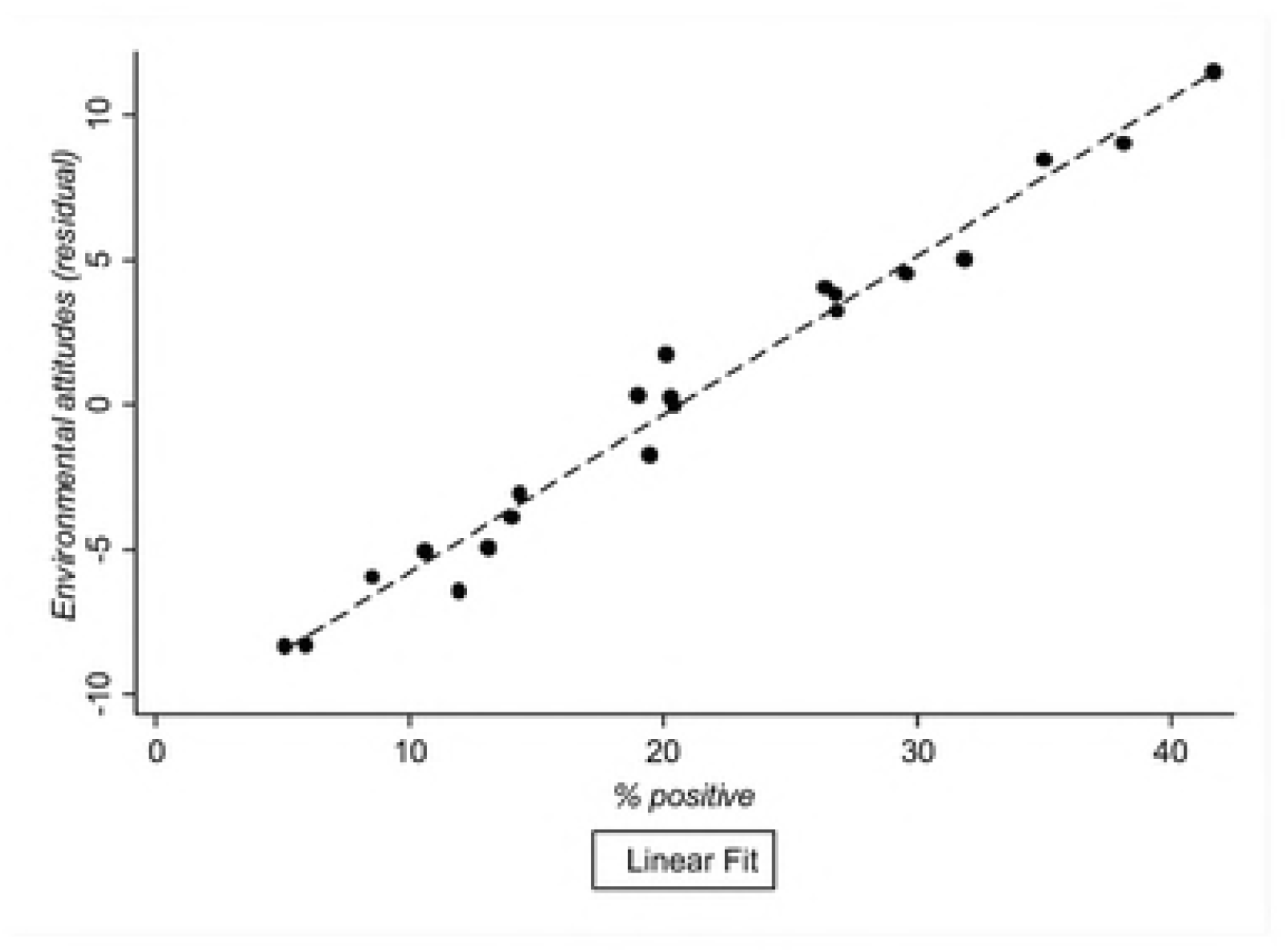
**Relationship between % positive and environmental attitudes in Canada.** *Note*. The y-axis represents the residualized value of environmental attitudes, which was computed by subtracting from the observed value, the value predicted with the estimated regression equation while holding % positive at its mean, i.e., predicted value = Observed Environmental Attitudes – [5041 -.37Year10 -.36 %negative + .61 (mean %positive) - 999.2Ln GDP + 49.27Ln GDP]^2^ (see Table 4).

**Figure 6.**
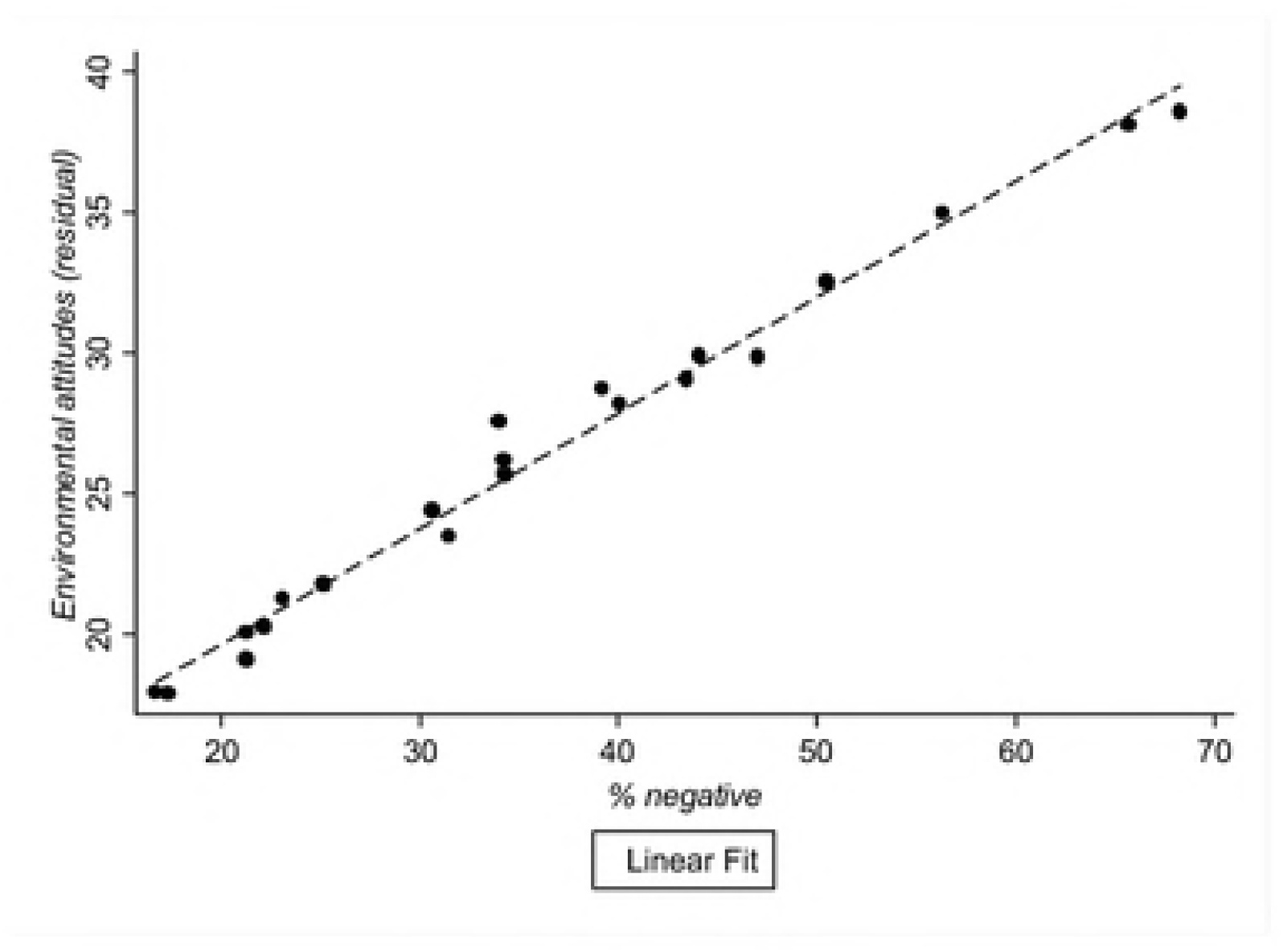
**Relationship between % negative and environmental attitudes in Canada** *Note.* The y-axis represents the residualized value of environmental attitudes, which was computed by subtracting from the observed value the value predicted with the estimated regression equation while holding % negative at its mean, i.e., predicted value = Observed Environmental Attitudes – [5041 -.37Year10 -.36 (mean %negative) + .61 %positive - 999.2Ln GDP + 49.27Ln GDP]^2^ (see Table 4).

#### Predicting subjective economy

Table 5 summarizes the results for the prediction of perceptions of the economy in Canada. Positive perceptions of the economy are significantly predicted by Equations 3 and 4, explaining 51% and 48% of the variance respectively (columns 1 and 2 in Table 5). The Year10 dummy did not reach significance, indicating that the historical trend prior to 2001 did not differ significantly from the historical trend thereafter as far as economic perceptions. Negative perceptions of the economy were also significantly related to the objective economy (see columns 3 and 4 in Table 5), explaining 75% of the variance for the model without a time lag (Equation 3) and 71% for the model with a time lag (Equation 4). Models without a time lag appear to do somewhat better overall.

The nonlinear trend shows a hump-shaped relationship between the objective economy and positive perceptions of the economy and a U-shaped relationship between the objective economy and negative perceptions (Figures 7 and 8). The inflection point was approximately $US27,000 GDP per capita (2011 price level). At lower levels in GDP per capita, positive perceptions are positively and negative perceptions are negatively related to the objective indicator as expected. However, similarly to Australia, this trend did not continue at higher levels of GDP per capita, beyond the inflection point. Again, the inflection point appears to act as a kind of reference point (Kahneman & Tversky, 1979), beyond which an improvement in the objective national economy does not contribute to its citizens’ economic sentiments.

**Table 5.**
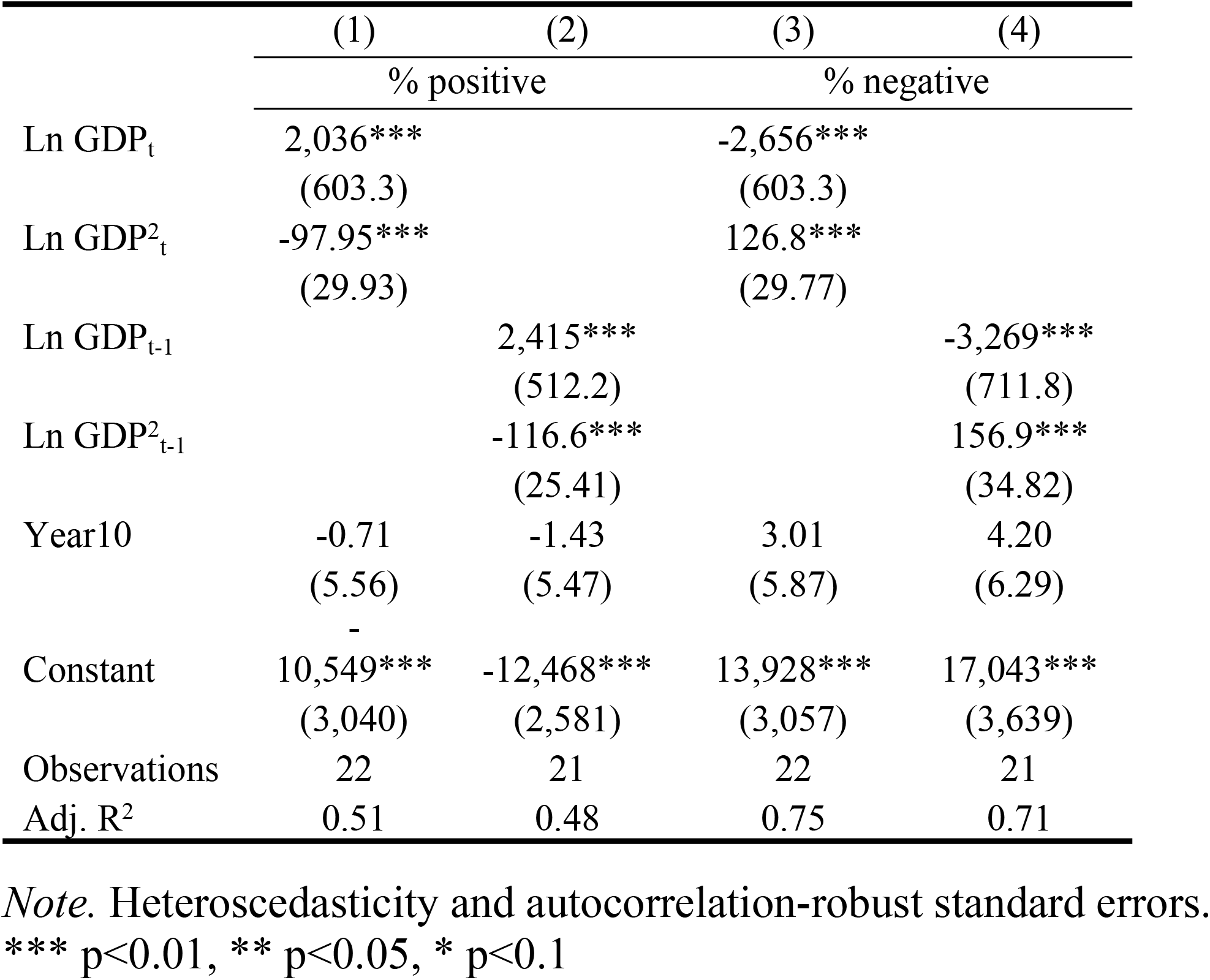
**Predicting subjective economy by objective economy in Canada.**

**Figure 7.**
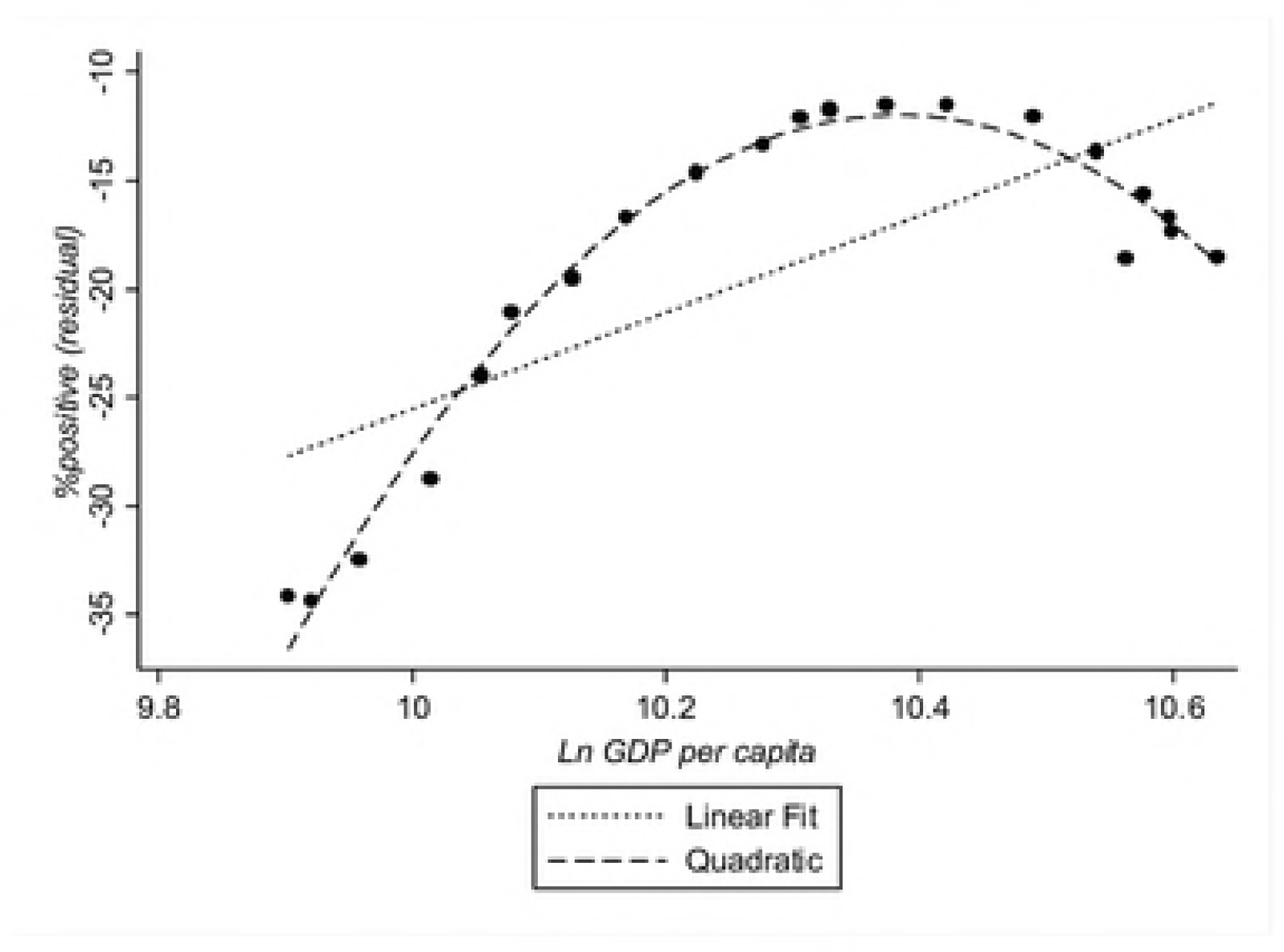
**Ln GDP per capita and % positive in Canada.** *Note.* The y-axis represents the residualized value of the proportion of the respondents who replied that the economy has gotten better (% positives), which was computed by subtracting from the observed value, the value predicted with the estimated regression equation while holding % positive at its mean, i.e., predicted value = Observed Environmental Attitudes – [13928 -3.01*year10 - 2656*(mean Ln GDP) + 126.8*(mean Ln GDP^2^] (see Table 5).

**Figure 8.**
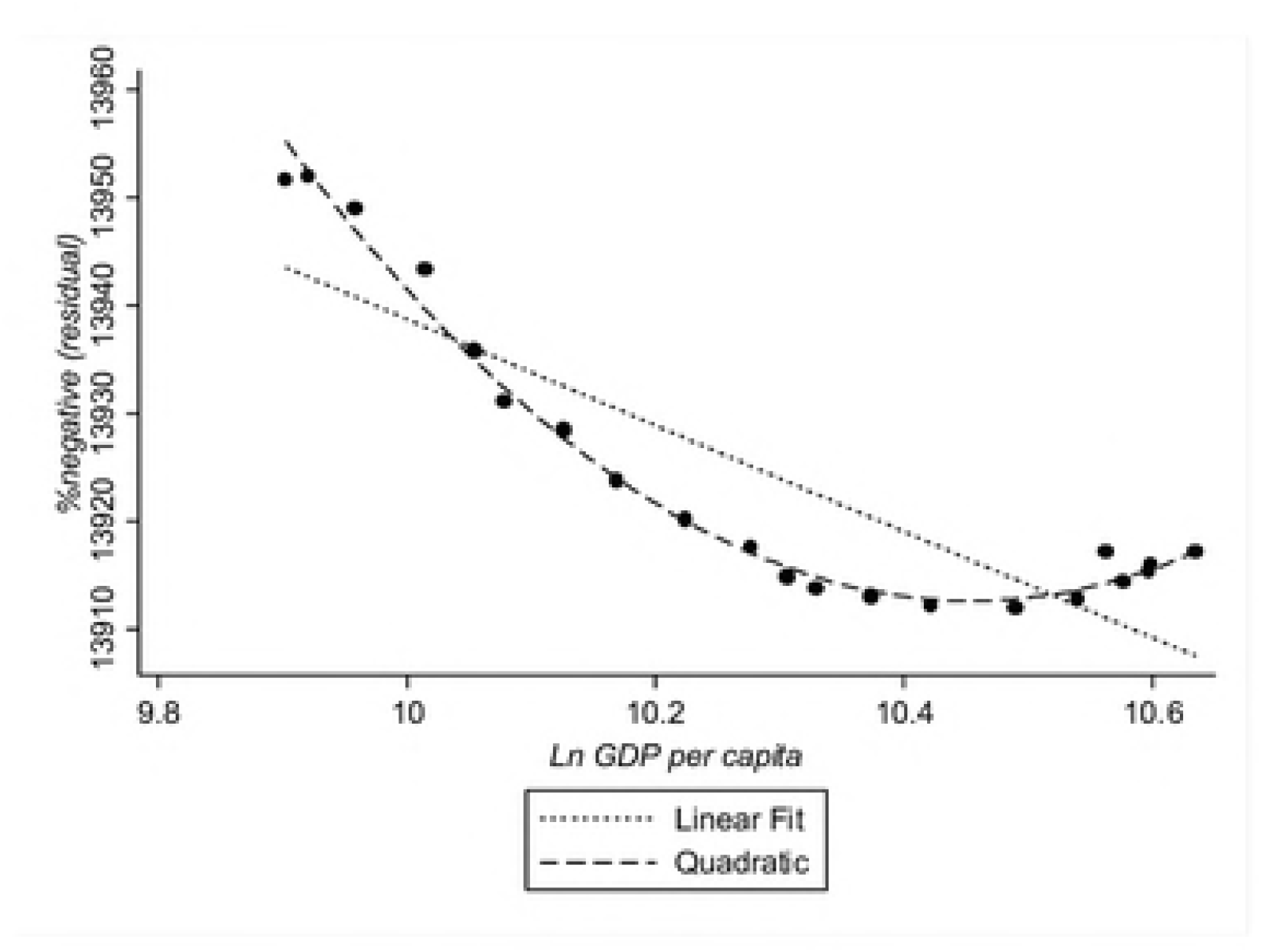
**Ln GDP per capita and % negative in Canada.** *Note.* The y-axis represents the residualized value of the proportion of the respondents who replied that the economy has gotten better (% positives), which was computed by subtracting from the observed value, the value predicted with the estimated regression equation while holding % positive at its mean, i.e., predicted value = Observed Environmental Attitudes – [10549 -0.71*year10 - 2036*(mean Ln GDP) + 97.95*(mean Ln GDP^2^] (see Table 5).

## Discussion

As the common wisdom has it, the economic condition of a nation is critically linked to the perceived importance of environment issues. However, there is a significant complexity to the relationship between the economy and environmental attitudes. First of all, there is an important commonality between Australia and Canada. In both countries, pro-environmental attitudes in a given year (i.e., proportion of people who regard the environmental issues as one of the most significant in time *t*) are well-predicted and correlated in a U-shaped fashion with GDP per capita of the same year (i.e., time *t*) or the preceding year (i.e., time *t-1*). In both countries, the fit of the models with and without a time lag was comparable; a minor difference between Australia and Canada may be due to slight differences in the wording of the survey items used (see Table 1). The shape of nonlinear trends was again similar. In the higher spectrum of GDP per capita, the better was the objective economic condition, the more environmental attitudes increased; in the lower spectrum of GDP per capita, the trend was reversed where the worse was the objective economic condition, the lesser were environmental attitudes. The inflection point was approximately $US30,000 and $US27,000 for Australia and Canada respectively. Second, the inclusion of perceptions of the economy significantly improved the prediction of environmental attitudes in both countries; positive perceptions of the economy increased the importance given to the environment. Third, the relationship between the objective economy and negative perceptions of the economy was again nonlinear; the relationship was negative in a lower range of objective economy, but was relatively flat in a higher range. The inflection point was again around US$30,000 (Australia) and US$27,000 (Canada) GDP per capita.

Nevertheless, there were subtle differences between Australia and Canada. First, in Canada, negative perceptions of the economy also predicted environmental attitudes over and above the nonlinear effects of the objective state of the economy and positive perceptions of the economy. As negative economic sentiments increased, environmental attitudes also increased. In Australia, however, this trend was present when only perceptions of the economy were included in the prediction of pro-environmental support (column 3 in Table 2); it became non-significant when objective economic indices were included in the model (columns 4 and 5, Table 2). Second, in Canada, both positive and negative perceptions of the economy were predicted by the objective indicator of the economy; however, in Australia, only negative perceptions of the economy were significantly predicted by the objective indicator. In combination, it appears that, in Australia, positive economic sentiments are driven by factors other than the nation’s overall economic performance, and therefore carries information other than the objective state of the economy. In addition, it is the positive economic sentiments that added significantly to the prediction of environmental attitudes in Australia. Future research should investigate further the determinants of environmental attitudes and positive and negative perceptions of a country’s economy in a larger sample of countries.

Let us present a conjecture here by way of providing a concise summary of the results. There appears to be a *reference point* on the dimension of national economy at which a nation’s citizenry shows a *baseline level of environmental attitudes*. This point may be regarded as a kind of national economic reference point, which its citizens expect as a satisfactory level of economic performance (as shown in the analyses of perceptions of the economy), so that an improvement beyond it is regarded as a gain, but a decline below it is seen as a loss (Kahneman & Tversky, 1979). Presumably, economic reference points and baseline environmental attitudes vary from one country to another—just as they vary across Australia and Canada. Nonetheless, as a nation’s economic performance departs from its reference point and its citizens’ perceptions of the economy are adjusted, environmental attitudes increase. On the one hand, when the economy is performing above the reference point and citizens perceive improving economic conditions, more citizens are positively disposed towards sustaining the natural environment. This is perhaps because a wealthy citizenry can afford to be concerned about the natural environment (Franzen & Meyer, 2010) and therefore are prepared to undertake pro-environmental actions (Inglehart, 1990, 1995). On the other hand, when the economy performs below the reference point and more citizens perceive worsening economic conditions, more citizens become increasingly concerned about the environment. We offer two speculations for this. First, it may be due to the national policy discourse and belief in ecological modernization (Hayden, 2014; Davidson & MacKendrick, 2004). When the economy is performing poorly, its growth can be spurred by boosting the investment in the green sectors of the industry, such as the development and uptake of renewable energies (see also Harring, Jagers, & Martinsson, 2011). A second possibility, found amongst Australian respondents in other studies, is a generalized feeling of societal pessimism or negative Zeitgeist (van der Bles, Postmes, & Meijer, 2015). As the economy shows a downturn and more citizens experience adverse economic circumstances, a pessimistic outlook may generalize to other domains of life as well; that is, the economy is bad, and therefore, so is the environment.

In summary, swings in economy relative to a reference point can raise different sentiments in the citizenry, thus affecting attitudes towards the environment. This conjecture needs to be more rigorously tested in future research.

## Conclusion

Objective indicators as well as subjective perceptions of the economic condition are significant influences on national environmental attitudes. At least in relatively wealthy countries such as Australia and Canada, the economy shows a U-shaped relationship with and environmental attitudes, which depends on the current state of economic performance. As the economic condition departs from some reference point, environmental attitudes become more prevalent. In contrast, when the economic performance falls below the reference point, the economic condition appears to have a negative relationship with environmental attitudes – the worse is a nation’s economy, the more environmentally concerned are its citizens. In addition, the *perceptions of the economy* can play a significant role, over and above those of objective indicators of the economy. Investigations of a larger sample of countries which incorporate both nonlinear trends of the objective economy and people’s subjective perceptions of the economy, are likely to provide greater depth in our understanding of the economic impact on environmental issues.

## Acknowledgments

This research was facilitated by a grant from the Australian Research Council to YK (DP130102229). We thank our colleagues at The University of Melbourne, Paul Dudgeon, Simon M. Laham, Léan V. O’Brien, Ilona M. McNeill, Rebecca C. Anderson, and Julia Meis, for their insight and expertise which greatly assisted the research.

## Supporting information

**S1 Dataset. Dataset – Australia.**

**S2 Dataset. Dataset – Canada.**

